# Phosphatidic acid inhibits inositol synthesis by inducing nuclear translocation of IP6K1 and repression of *myo*-inositol-3-P synthase

**DOI:** 10.1101/2022.02.21.481348

**Authors:** Pablo Lazcano, Michael W. Schmidtke, Chisom Onu, Miriam L. Greenberg

## Abstract

Inositol is an essential metabolite that serves as a precursor for structural and signaling molecules. Although perturbation of inositol homeostasis has been implicated in numerous human disorders, surprisingly little is known about how inositol levels are regulated in mammalian cells. A recent study in mouse embryonic fibroblasts (MEFs) demonstrated that nuclear translocation of inositol hexakisphosphate kinase 1 (IP6K1) mediates repression of *myo-*3-P synthase (MIPS), the rate-limiting inositol biosynthetic enzyme. Binding of IP6K1 to phosphatidic acid (PA) is required for this repression. The current study was carried out to elucidate the role of PA in IP6K1 repression. The results indicate that increasing PA levels through pharmacological stimulation of phospholipase D (PLD) or direct supplementation of 18:1 PA induces nuclear translocation of IP6K1 and represses expression of MIPS protein. This effect was specific to PA synthesized in the plasma membrane, as ER-derived PA did not induce IP6K1 translocation. PLD-mediated PA synthesis can be stimulated by the master metabolic regulator 5’ AMP-activated protein kinase (AMPK). Activation of AMPK by glucose deprivation or by treatment with the mood stabilizing drugs valproate (VPA) or lithium recapitulated IP6K1 nuclear translocation and decreased MIPS expression. This study demonstrates for the first time that modulation of PA levels through the AMPK-PLD pathway regulates IP6K1-mediated repression of MIPS.

## INTRODUCTION

Inositol is a six-carbon cyclitol that is essential for all eukaryotic life (Case et al., 2020). Deprivation of inositol results in “inositol-less death”, a phenotype originally described in yeast and supported by early work in human cell lines (Eagle et al., 1957; Keith et al., 1977; Matile, 1966; Minskoff et al., 1992). The importance of inositol is due in part to its role as a precursor for other vital molecules, such as phosphoinositides, inositol polyphosphates, and pyrophosphates (Case et al., 2020). Cells must obtain inositol through extracellular uptake via inositol transporters or *de novo* synthesis from glucose-6-phosphate (G6P).

Dysregulation of inositol metabolism has been linked to several prevalent human disorders, including Alzheimer’s, polycystic ovary syndrome, Lowe syndrome, various cancers, and bipolar disorder (Artini et al., 2013; Kassie et al., 2010; Papaleo et al., 2007; Shi et al., 2006). Furthermore, inositol depletion has been hypothesized as the mechanism of action of the mood stabilizing drugs lithium and valproate (VPA), as both are known to inhibit the inositol biosynthetic pathway and thereby decrease inositol levels (Berridge, 2014; Shaltiel et al., 2004; Vaden et al., 2001; Williams et al., 2002). Despite its critical importance, little is known about the mechanisms that regulate inositol biosynthesis in mammalian cells.

Some of the first insights regarding regulation of inositol synthesis in mammalian cells came from a study showing that inositol hexakisphosphate kinase 1 (IP6K1) negatively regulates expression of the rate-limiting enzyme in the inositol biosynthetic pathway, *myo*-inositol-3-phosphate synthase (MIPS). Accordingly, IP6K1 repressed MIPS expression by increasing DNA methylation in the promoter of its encoding gene, ISYNA1 (Yu et al., 2016). The authors determined that IP6K1-mediated repression of MIPS requires binding of IP6K1 to phosphatidic acid (PA).

PA can be generated through three distinct cellular pathways. These include (1) acylation of lyso-PA by lyso-PA acyltransferase, (2) phosphorylation of diacylglycerol by diacylglycerol kinase, and (3) hydrolysis of phosphatidylcholine (PC) by phospholipase D (PLD) (Foster et al., 2014; Kassas et al., 2017; Liscovitch et al., 2000; Thakur et al., 2019). PLD activity is regulated through phosphorylation by the highly conserved kinase AMPK (Kim et al., 2010). As a master metabolic regulator, canonical activation of AMPK is mediated by an increase in the AMP/ADP:ATP ratio (Garcia and Shaw, 2017). Interestingly, the inositol-depleting drugs VPA and lithium have been shown to activate AMPK in multiple cell types, and VPA was shown to activate the AMPK homolog Snf1 in yeast (Avery and Bumpus, 2014; Bao et al., 2019; Lee et al., 2020; Salsaa et al., 2021). Furthermore, a recent study identified inositol as a direct allosteric inhibitor of AMPK that acts by blocking activation by AMP (Hsu et al., 2021). Taken together, these reports suggest a potential feedback mechanism involving PA, AMPK, IP6K1, and inositol synthesis.

Based on the above findings, we hypothesized that modulation of PA levels through the AMPK-PLD pathway serves as the regulatory mechanism underlying repression of inositol biosynthesis by IP6K1. In the current study, increasing PA levels by drug-mediated stimulation of PLD or activation of AMPK resulted in increased nuclear translocation of IP6K1 and decreased expression of MIPS. These findings delineate the first mechanistic model of regulation of inositol biosynthesis in mammalian cells in which PA controls IP6K1-mediated repression of MIPS by increasing IP6K1 localization in the nucleus.

## EXPERIMENTAL PROCEDURES

### Cell culture, transfections, and treatments

WT and IP6K1-KO MEF cells (Yu et al., 2016) were cultured in DMEM (Gibco) supplemented with 10% FBS (R&D Systems) and 1% penicillin-streptomycin (Gibco). For imaging, cells were grown either on glass coverslips or glass-bottom 35 mm dishes (MatTek). Cells grown on coverslips were washed with PBS and fixed with a 4% solution of paraformaldehyde/sucrose in PBS. The coverslips were mounted onto microscope slides using ProLong Diamond Antifade Mountant with DAPI (Invitrogen). Cells grown in glass-bottom dishes were imaged without fixation. Constructs of GFP-PASS and RFP-PASS (Zhang et al., 2014) were a kind gift from the laboratory of Dr. Guangwei Du at the University of Texas Health Science Center at Houston. Plasmids for optogenetic induction of PLD: optoPLD-PM (Addgene plasmid # 140114), optoPLD-PM ‘dead’ (Addgene plasmid # 140061), optoPLD-ER (Addgene plasmid # 140060) and optoPLD-ER ‘dead’ (Addgene plasmid # 140059) (Tei and Baskin, 2020) were constructed by the laboratory of Dr. Jeremy Baskin from Cornell University, Ithaca, NY, and purchased through Addgene. 500-1000 ng of plasmid was transfected using Lipofectamine 3000 transfection reagent (Thermo Fisher Scientific) following the manufacturer’s instructions. For co-transfections of two plasmids, a ratio of 1:1 was used. Transfection complexes were prepared in Opti-MEM (Thermo Fisher Scientific). All experiments were performed 24 – 48 h after transfection to ensure proper expression of the plasmids. Phorbol 12-myristate 13-acetate (PMA) (Sigma-Aldrich, P8139), FIPI hydrochloride hydrate (Sigma-Aldrich, F5807), 18:1 Phosphatidic Acid (Avanti, 840875), Lithium Chloride (Sigma-Aldrich, L9650) and Valproic Acid (Cayman, 13033) were used for cell treatments. Concentration and times of treatment are included in the corresponding figure legends.

### Optogenetic induction of PA synthesis

A blue light box was constructed with blue LED strips (1000bulbs.com- FLX-00036) surrounding an acrylic box with a lid. The LED strips were taped on the outside of the box with the lightbulbs facing in. The box was completely covered in black felt paper. The system was connected to a remote control. The box was placed inside a CO_2_ incubator. Plates of cells transfected with optoPLD plasmids were placed inside the light box and the incubator door was shut to allow for normal CO_2_ and humidity conditions. Cells inside the light box were stimulated with remotely controlled intermittent blue light in pulses of 5 seconds every 2 minutes, for a total of 20-30 minutes. After stimulation, cells were analyzed immediately by confocal microscopy.

### Protein extraction and Western blotting

Total cell lysates were obtained using a radioimmunoprecipitation assay (RIPA) buffer (Santa Cruz Biotechnology). Protein concentration was determined using a Bradford protein assay (Bio-Rad) against a bovine serum albumin standard. 30-50 μg of total protein per sample was run on a 12% SDS-PAGE gel and transferred to a polyvinylidene difluoride (PVDF) membrane. Primary antibodies against MIPS (Invitrogen #PA5-44105, 1:500 dilution), p-AMPK (Cell Signaling #2531S, 1:1000 dilution), AMPK (Cell Signaling #2793S, 1:1000 dilution), IP6K1 (GeneTex # GTX103949, 1:5000 dilution) and Actin (Santa Cruz Biotechnology #, 1:20000 dilution) were used. Corresponding europium-tagged secondary antibodies (Molecular Devices) were used at a 1:5000 dilution and 1:20,000 for actin, and the signal was detected using a SpectraMax i3x plate reader and ScanLater Western blotting cartridge (Molecular Devices). ImageJ was used for band intensity quantification. Actin was used to normalize for protein loading.

### Image analysis

All images were obtained on a Leica SP8 confocal microscope. Fluorescence quantification, analysis, and post-acquisition detailing of images was performed with ImageJ.

### Statistical analysis

All graphs show average values ± SD. Statistical analyses were performed with GraphPad Prism software, version 6.1 (GraphPad, San Diego,CA). For column graphs, significant differences between experiments were assessed by an unpaired t-test or a one-way ANOVA with a Tukey posttest, considering α as 0.05. For fluorescence graphs, significant differences between experiments were assessed by a Kolmogorov-Smirnov test, considering α as 0.05.

## RESULTS

### PA regulates IP6K1 localization

A previous study demonstrated that nuclear translocation of IP6K1 requires PA binding and that mRNA levels of the inositol biosynthetic enzyme MIPS are reduced in IP6K1-KO MEF cells (Yu et al., 2016). To delineate the role of PA, we tested the hypothesis that modifying PA levels regulates MIPS expression by controlling IP6K1 localization. To modify PA levels, we targeted the PLD-mediated PA synthesis pathway, as PLD activity is amenable to pharmacological treatment and is endogenously regulated by AMPK, a putative target of inositol-depleting mood stabilizers (Hsu et al., 2021; Kim et al., 2010). MEF cells were treated with either PMA, a drug known to induce PA synthesis through PKC-dependent activation of PLD (Hu and Exton, 2003; Liu et al., 2017), or FIPI, a potent PLD1/2 inhibitor (Ganesan et al., 2015). To confirm the efficacy of these treatments for modifying PA levels, the genetically-encoded PA biosensors RFP-PASS and GFP-PASS were used to visually monitor changes in PA abundance (Zhang et al., 2014). These constructs contain a strong PA-binding domain linked to either RFP or GFP as a fluorescent marker. MEF cells transfected with RFP-PASS were treated with 100 nM PMA for 5h in the presence or absence of 0.75 μM FIPI and analyzed by confocal microscopy. As expected, PMA treatment induced PA synthesis, as evidenced by an approximately 2-fold increase in RFP fluorescence at the plasma membrane (Figure 1). Pretreatment with FIPI blocked the increase in RFP fluorescence, confirming that the effect of PMA on PA levels is PLD-dependent (Figure 1).

**FIGURE 1.**
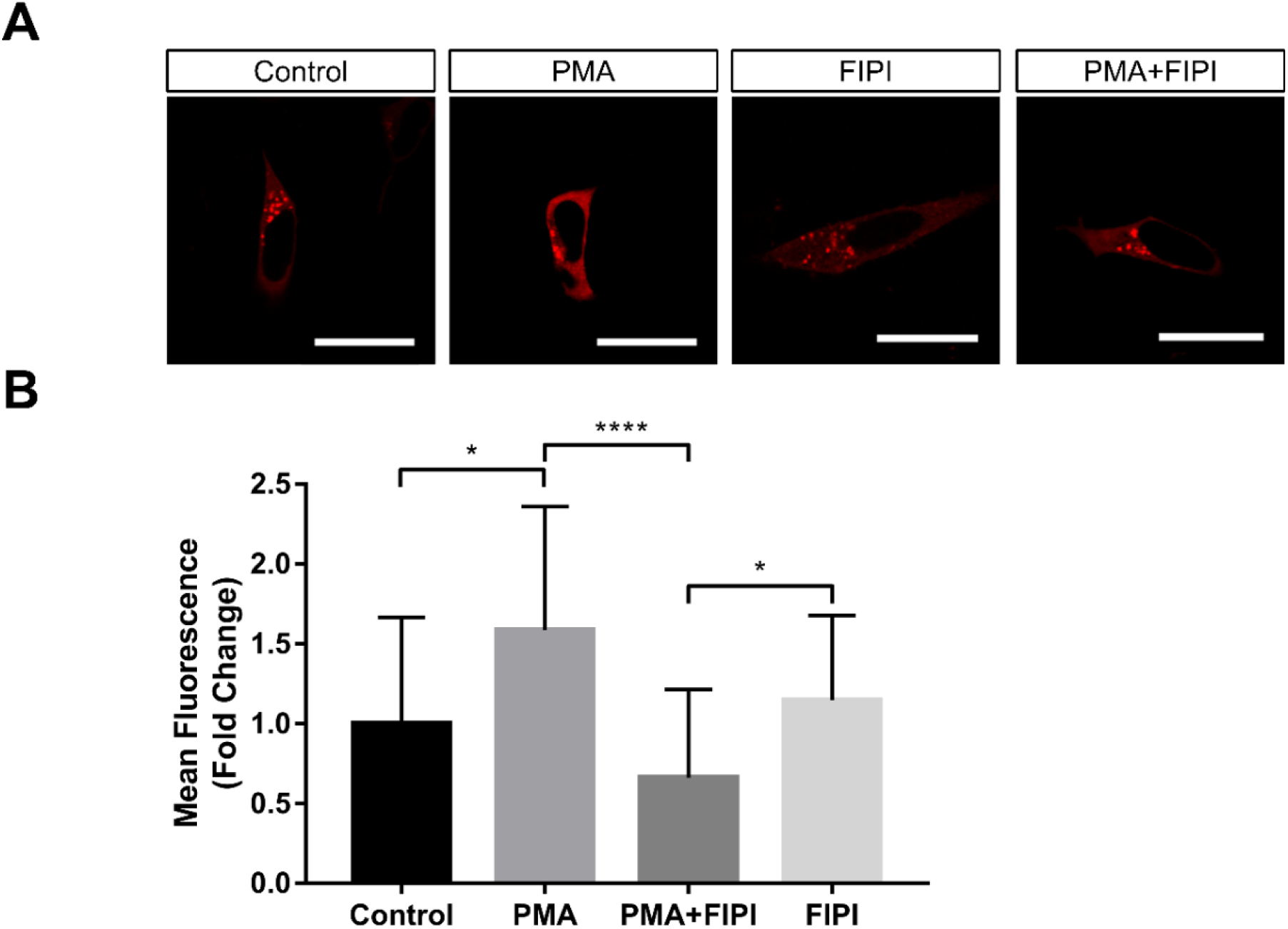
PA levels are modulated by treatment with PMA or FIPI. A, Cells transfected with RFP-PASS (red fluorescence) were treated with PMA alone (100 nM for 5h) or in the presence of FIPI (0.75 μM, applied 30 min before PMA). n = 3 independent experiments, images are representative of the observed effect in most cells. Scale bars = 20 μm. B, Mean fluorescence quantification of experiment depicted in ‘A’. Arbitrary fluorescence units were normalized to untreated (control) cells and expressed as fold change ± SD. Statistical significance was analyzed by one-way ANOVA with a Tukey *post hoc* test, *p<0.05, ****p<0.0001 where n = 3 independent experiments in which 20-30 cells were analyzed per condition.

To track IP6K1 localization, IP6K1-KO MEF cells were transfected with an IP6K1-GFP expression plasmid (Yu et al., 2016). This approach ensured that all IP6K1 in the cell could be visualized by GFP fluorescence. To modify PA levels, cells were treated with PMA or cell-permeable 18:1 PA (Hatton et al., 2015) in the presence or absence of FIPI, then localization of IP6K1 was monitored by confocal microscopy. IP6K1 is normally localized in both the cytosol and nucleus; however, following treatment with either PMA or 18:1 PA, nuclear localization of IP6K1 was notably increased (Figure 2a and 2b). As expected, pretreatment with FIPI prevented the change in IP6K1 localization in PMA-treated cells but not in cells supplemented with exogenous 18:1 PA, which accumulate PA independent of PLD activity (Figure 2a and 2b). Total IP6K1 protein levels were unaffected by either treatment (Figure 2c). These results demonstrate that MEF cells accumulate IP6K1 in the nucleus as a function of PA levels.

**FIGURE 2.**
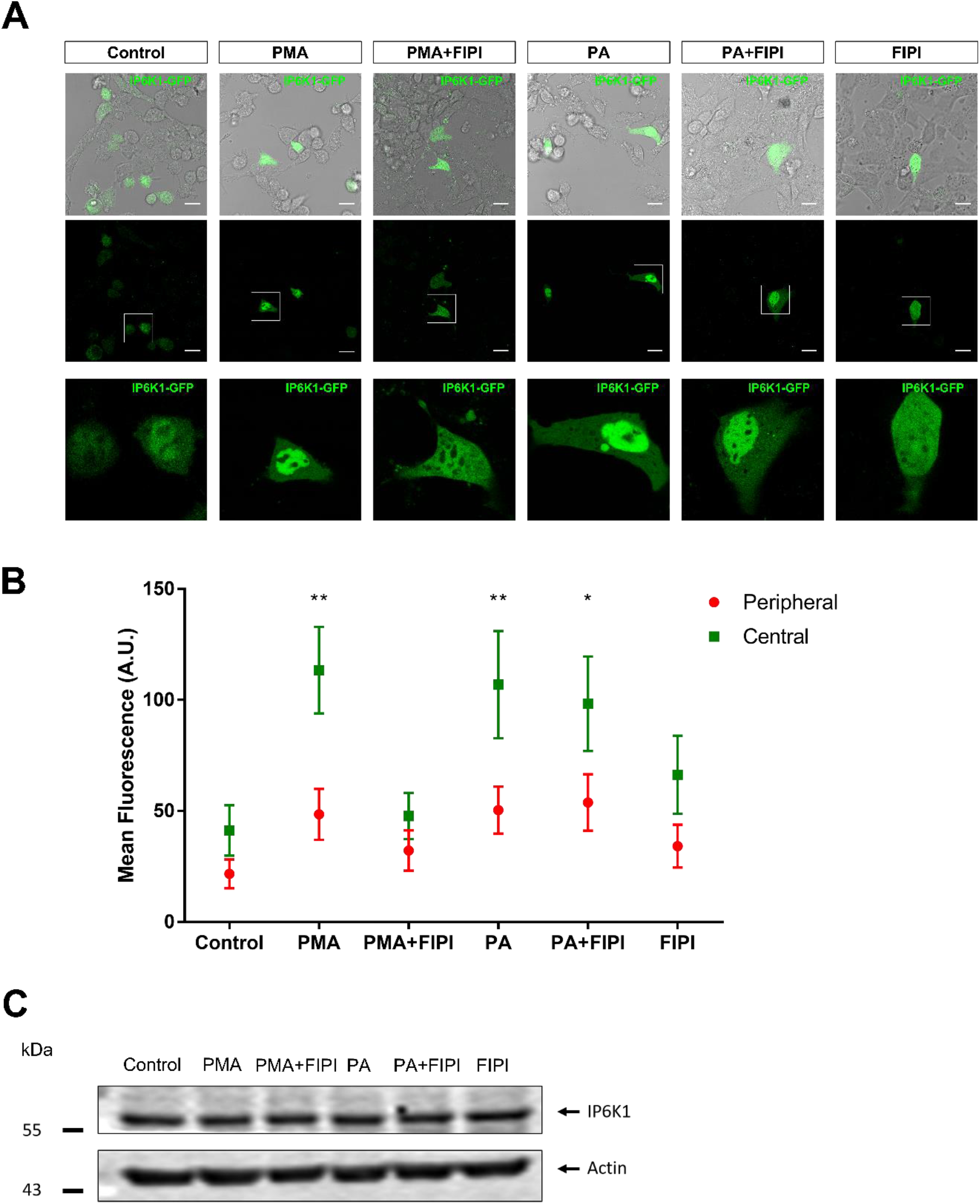
IP6K1 localization is regulated by PA. A, IP6K1-KO cells transfected with IP6K1-GFP (green) were treated with 100 nM PMA for 16h or 100 μM 18:1 PA for 5 h, in the presence or absence of FIPI (0.75 μM, applied 30 min before PMA or PA). Top panel: merged image of transmitted light with green fluorescence image to evaluate transfection efficiency. Middle panel: green fluorescence alone; boxed region is magnified to show further detail in the lower panel. Scale bars = 20 μm. B, Mean fluorescence (arbitrary units, A.U.) of peripheral (cytosolic) and central (nuclear) sections of cells as shown in A. Statistical significance was analyzed by a Kolmogorov-Smirnov test, *p<0.05, **p<0.01 where n = 3 independent experiments in which 20-30 cells were analyzed per condition. C, Representative Western blot analysis of IP6K1 in WT MEF cells treated as in A. n = 3 independent experiments, image is representative of the effect observed in all experiments. Actin was used to normalize for protein loading.

PA is synthesized in multiple subcellular locations, and the acyl chain composition of newly synthesized PA is influenced by the site of synthesis, reflecting differences in the composition of available precursor lipids from which PA is synthesized (Du et al., 2004; Fazio et al., 2020; Zhukovsky et al., 2019). The source of PA is critical for its regulatory functions. For example, it was recently demonstrated that only PA synthesized in the plasma membrane (but not the ER, Golgi, or endosomes) can regulate Hippo pathway signaling (Han et al., 2018; Tei and Baskin, 2020). Therefore, to test whether the source of PA affects its ability to induce IP6K1 translocation, an optogenetic approach (optoPLD) was adopted wherein PA synthesis by PLD can be targeted to specific subcellular membranes and temporally regulated by exposure to blue light (Tei and Baskin, 2020).

Using this approach, PA synthesized at the plasma membrane (PM-optoPLD) was compared with ER-derived PA (ER-optoPLD) for its ability to stimulate nuclear localization of IP6K1. Co-transfection of the PA biosensor GFP-PASS with either PM-optoPLD or ER-optoPLD confirmed proper targeting of PA synthesis to the PM and perinuclear ER, respectively (Figure 3a). This effect did not occur using the corresponding catalytically ‘dead’ control constructs. Interestingly, only PM-derived PA (but not ER-derived PA) was able to stimulate IP6K1 nuclear translocation (Figure 3b). As expected, nuclear localization was not stimulated by the catalytically ‘dead’ constructs. Collectively, these findings further support the hypothesis that PA regulates nuclear translocation of IP6K1 and indicate that PA synthesized by PLD in the PM is unique in its ability to mediate this effect.

**FIGURE 3.**
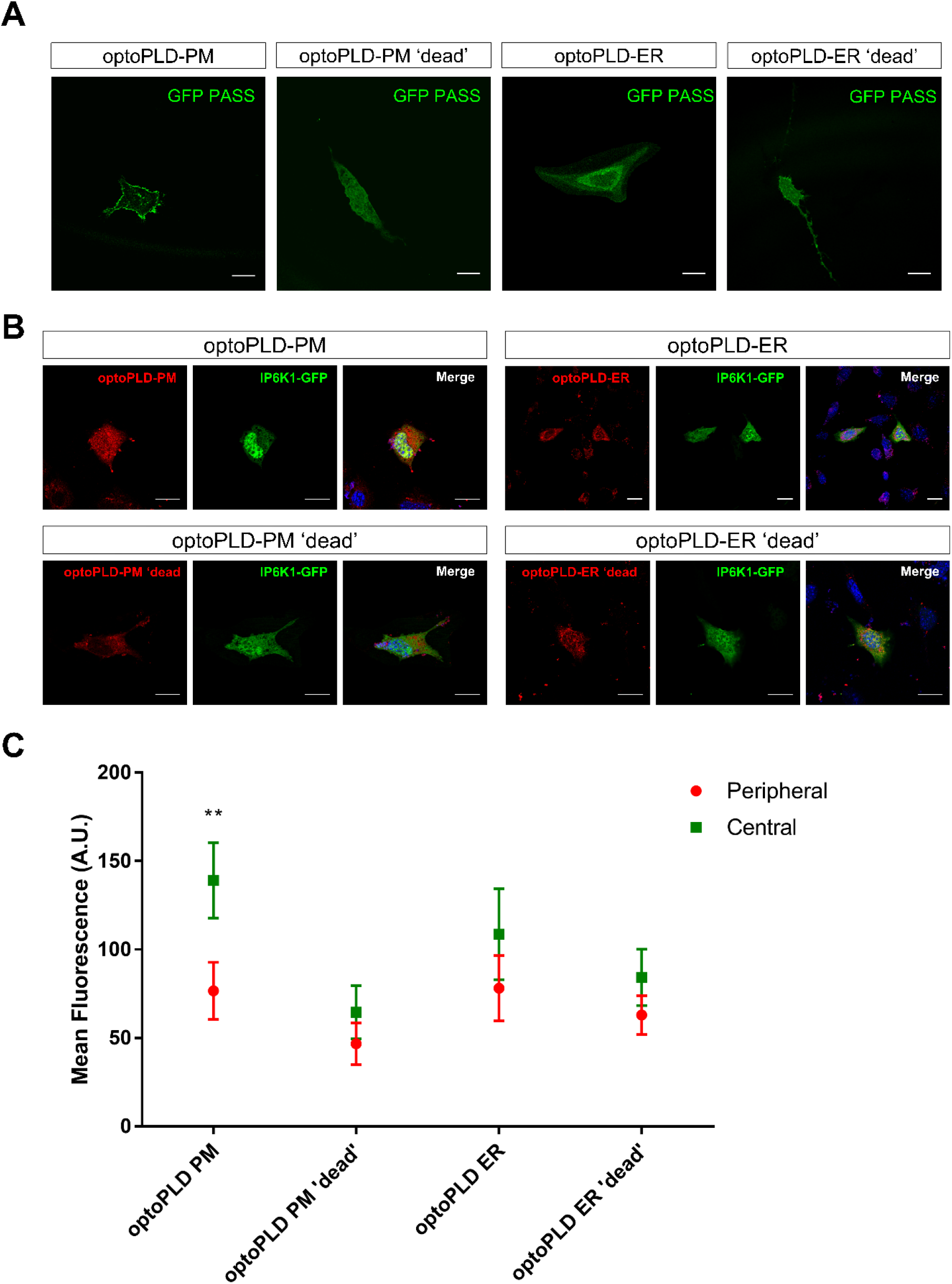
Plasma membrane-derived PA induces IP6K1 nuclear localization. A, MEF cells were co-transfected with GFP-PASS (green) and either optoPLD-PM, optoPLD-PM ‘dead’, optoPLD-ER, or optoPLD-ER ‘dead’ as indicated. Cells were stimulated using intermittent blue light with a 5-second pulse every 2 min for a total of 30 min. PA localization was evaluated by GFP-PASS fluorescence using confocal microscopy. Scale bar = 20 μm. Images are representative of the observed phenotype. B, IP6K1-KO MEF cells were co-transfected with IP6K1-GFP (green) and optoPLD-ER (red, top left), optoPLD-PM (red, top right), optoPLD-ER ‘dead’ (red, bottom left), or optoPLD-PM ‘dead’ (red, bottom right). Cells were stimulated using intermittent blue light with a 5-second pulse every 2 min for a total of 30 min. IP6K1 localization was determined by confocal microscopy. DAPI was used as nuclear staining. Scale bar = 20 μm. C, Mean fluorescence (arbitrary units, A.U.) of peripheral (cytosolic) and central (nuclear) sections of cells as shown in B. Statistical significance was analyzed by a Kolmogorov-Smirnov test, **p<0.01 where n = 3 independent experiments in which 20-30 cells were analyzed per condition.

### PA regulates MIPS expression

Yu *et al*. showed that IP6K1-KO cells have higher levels of MIPS mRNA and protein compared to WT MEF cells, suggesting that IP6K1 serves as a negative regulator of MIPS expression (Yu et al., 2016). To test the hypothesis that PA levels modulate MIPS expression by controlling IP6K1 localization, we treated cells with PMA in the presence or absence of FIPI and measured MIPS protein levels by Western blot. Treatment with either PMA or 18:1 PA resulted in a significant reduction in MIPS protein (Figure 4a), and as expected, only the PMA-induced decrease was blocked by pretreatment with FIPI (Figure 4a). Strikingly, regulation of MIPS protein levels by PA did not occur in IP6K1-KO MEF cells, indicating that IP6K1 is required for this regulatory mechanism to function (Figure 4b). Taken together, these results show that MIPS expression is regulated by PA levels in an IP6K1-dependent manner.

**FIGURE 4.**
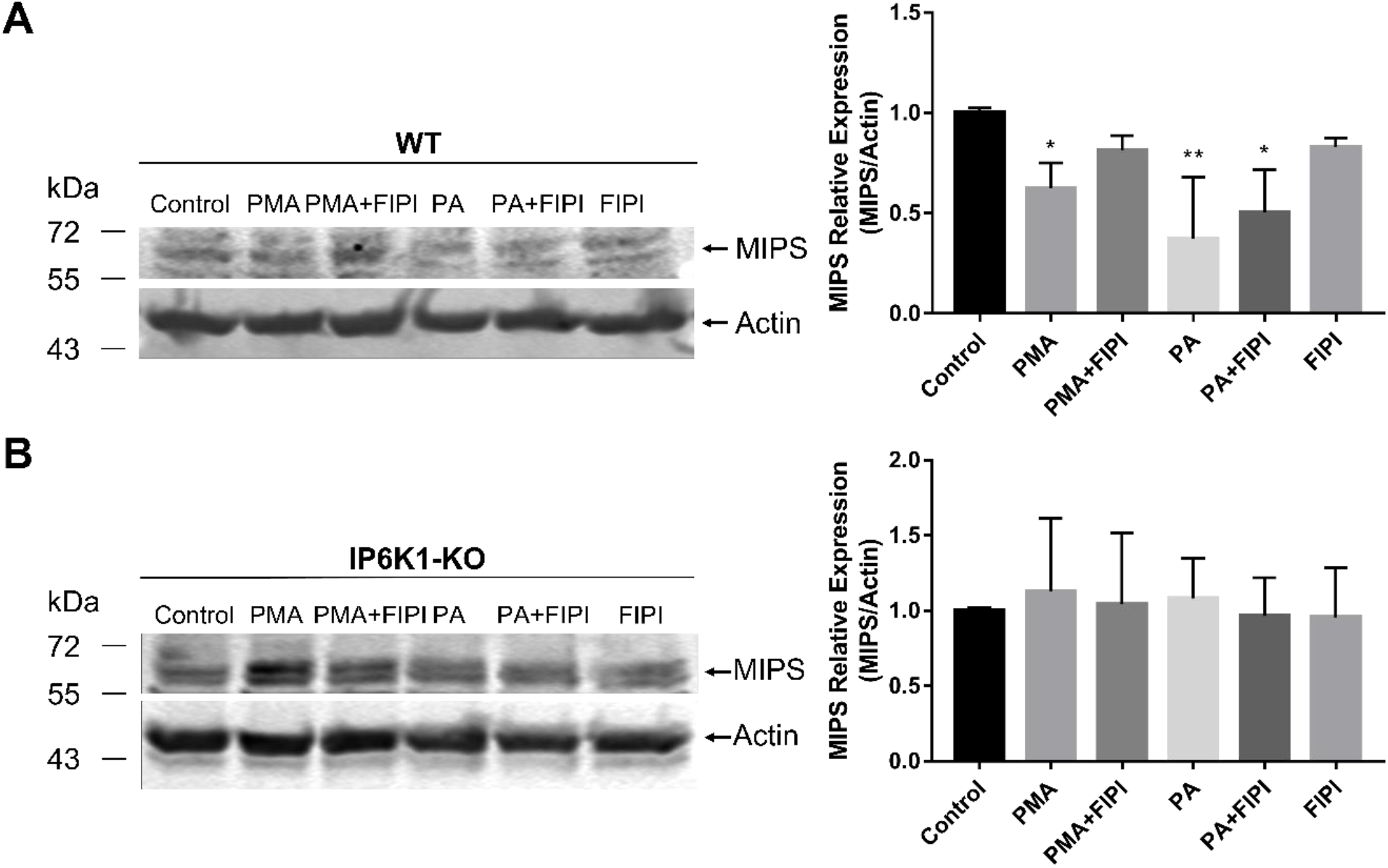
MIPS expression is regulated by PA in the presence of IP6K1. A, Western blot against MIPS protein in WT MEF cells treated with 100 nM PMA for 16 h or 100 μM 18:1 PA for 5h, in the presence or absence of FIPI (0.75 μM, applied 30 min before PMA or PA). B, Western blot against MIPS protein in IP6K1-KO MEF cells treated with 100 nM PMA for 16 h or 100 μM 18:1 PA for 5h, in the presence or absence of FIPI (0.75 μM, applied 30 min before PMA or PA). Graphs depict band quantification (right). Actin was used to normalize for protein loading. n = 3 independent experiments for each condition. Statistical significance was analyzed by one-way ANOVA with a Tukey *post hoc* test, *p<0.05 and **p<0.01.

### Increasing PA levels by glucose deprivation induces nuclear translocation of IP6K1 and MIPS repression

The above findings utilized drug treatments and exogenous supplementation of PA to alter intracellular PA levels. To complement these experiments, we took an orthogonal approach to assay nuclear translocation of IP6K1 under physiological conditions that stimulate PA synthesis. PLD1-mediated PA synthesis is increased in response to activation of the highly conserved kinase AMPK (Kim et al., 2010). AMPK is widely recognized as a master metabolic regulator that is activated by phosphorylation when energy is low to shift cells from anabolism to catabolism (Garcia and Shaw, 2017; Hardie et al., 2012). Numerous studies reported increased AMPK activation in response to glucose limitation (Endo et al., 2018; Lin and Hardie, 2018; Salt et al., 1998; Zhang et al., 2017). Therefore, to determine the effect of energy stress-induced PA synthesis on localization of IP6K1, cells were grown in glucose-free conditions for 16h and PA abundance and IP6K1 localization were monitored. As seen in Figure 5a and 5b, glucose-deprived cells exhibited a 3-fold increase in RFP-PASS fluorescence, indicative of increased PA synthesis. Increased PA correlated with an increase in the nuclear localization of IP6K1 and a marginally significant reduction in MIPS protein levels (p= 0.09; Figure 5c and 5d).

**FIGURE 5.**
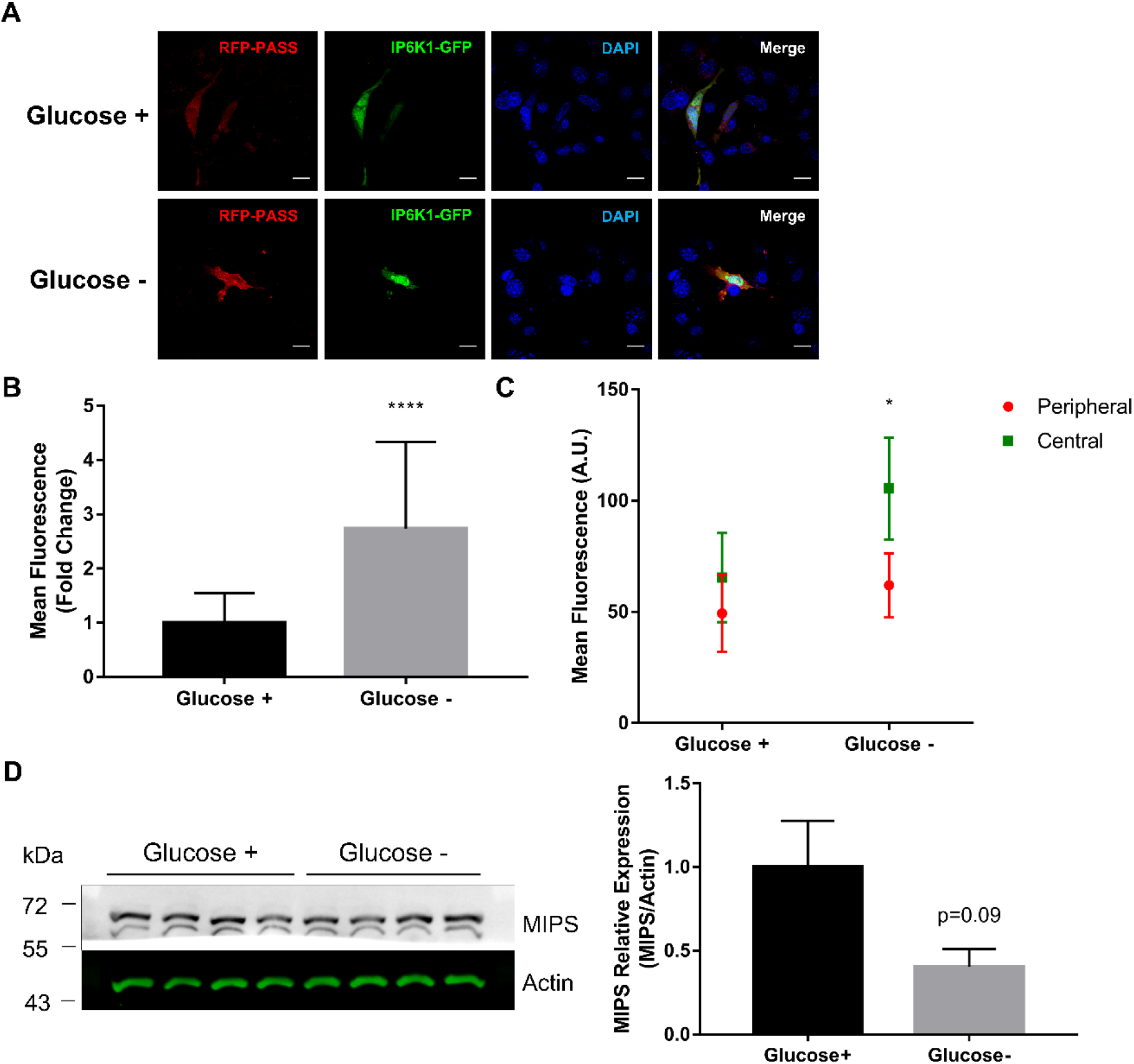
Glucose deprivation induces PA synthesis, IP6K1 nuclear translocation, and MIPS repression. A, IP6K1-KO MEF cells were co-transfected with IP6K1-GFP and RFP-PASS. Cells were deprived of glucose for 16 h and PA synthesis and IP6K1 localization were monitored by confocal microscopy. DAPI was used to stain nuclei. Scale bar = 20 μm. B, Mean fluorescence quantification of RFP-PASS from experiment depicted in A. Arbitrary fluorescence units (A.U.) were normalized to control and expressed as fold change ± SD. Statistical significance was analyzed by an unpaired t-test, ****p<0.0001, where n = 3 independent experiments and between 20-30 cells were analyzed per condition. C, mean fluorescence (arbitrary units, A.U.) of peripheral (cytosolic) and central (nuclear) sections of cells as shown in A. Statistical significance was analyzed by a Kolmogorov-Smirnov test, *p<0.05 where n = 3 independent experiments and between 20-30 cells were analyzed per condition. D, Western blot against MIPS protein in WT MEF cells grown in control conditions (glucose +) or deprived of glucose for 16h (glucose -). Graphs depict band quantification (right). Actin was used to normalize for protein loading. n = 4 independent experiments for each condition. Statistical significance was analyzed by an unpaired t-test.

### Activation of AMPK by lithium or VPA induces nuclear localization of IP6K1 and MIPS repression

The inositol-depleting mood stabilizing drugs lithium and VPA have been shown to activate AMPK (Avery and Bumpus, 2014; Bao et al., 2019; Salsaa et al., 2021). To test the prediction that activation of AMPK by mood stabilizers induces nuclear localization of IP6K1, cells were treated with VPA (1 mM) or lithium (10 mM) for 24 h, and phosphorylated (active) AMPK was measured by Western blot (Willows et al., 2017). Treatment with either VPA or lithium increased phosphorylation of AMPK by 50% and 100%, respectively, relative to controls (Figure 6a and 6b). The total amount of AMPK protein was not altered by either treatment. Localization of IP6K1-GFP in cells treated with lithium or VPA for 24 h was examined by confocal microscopy. Nuclear localization of IP6K1 was increased by treatment with lithium or VPA (p< 0.05 and p= 0.07 respectively; Figure 6c and 6d), and importantly, these treatments induced a 50% decrease in MIPS protein levels (p= 0.053 and p< 0.05 respectively; Figure 6e).

**FIGURE 6.**
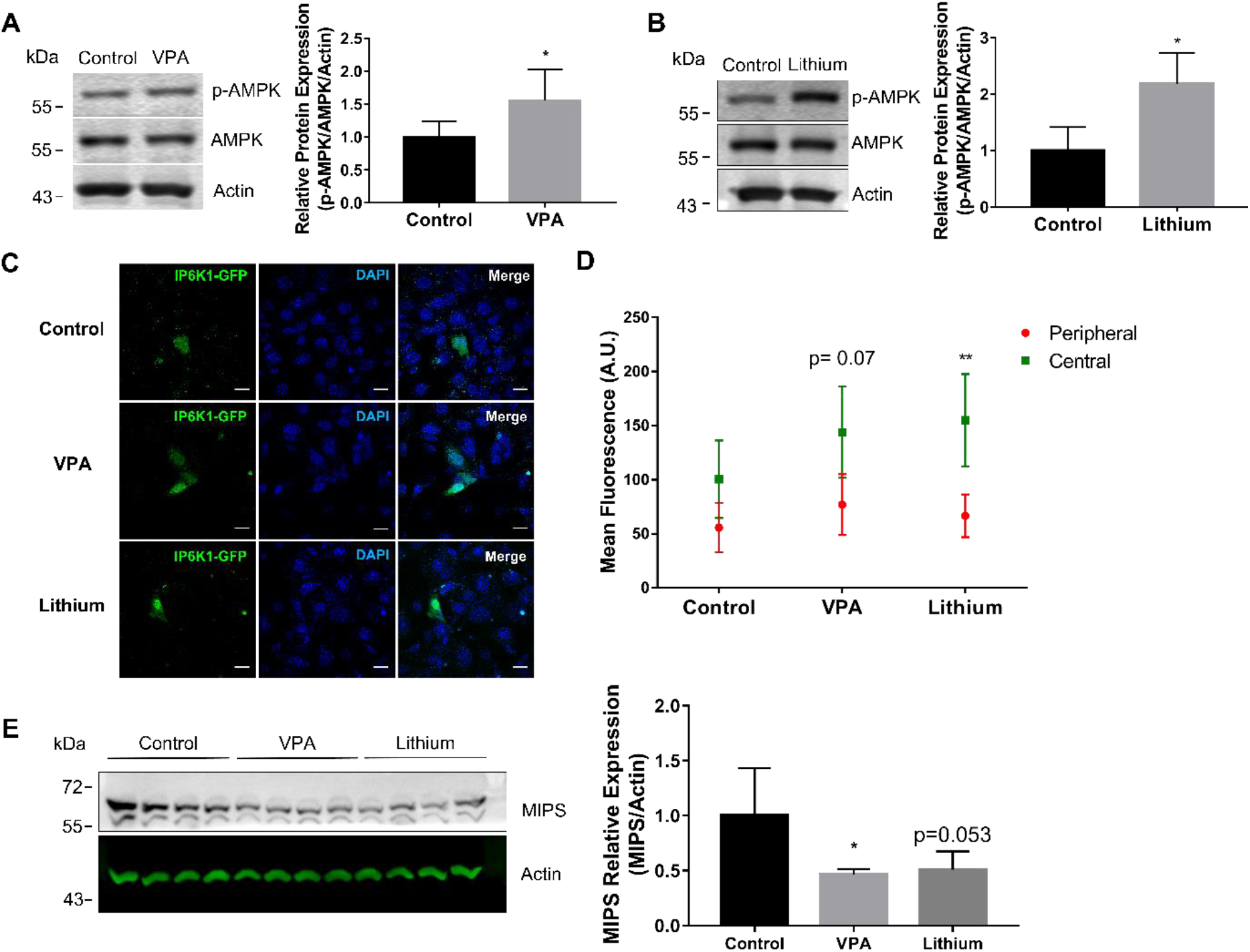
VPA and lithium activate AMPK, induce IP6K1 nuclear translocation, and repress MIPS protein expression. A, Cells were treated with 1 mM VPA for 24 h and AMPK activation was determined by Western blot. Samples were probed for p-AMPK, stripped until no signal was detected and re-probed for total AMPK. The lower part of the membrane was probed against actin to normalize for protein loading. The graph shows the relative protein expression of p-AMPK normalized against total AMPK and then actin. Statistical significance was analyzed by parametric t-test, *p<0.05 where n = 5 independent experiments. B, Cells were treated with 10 mM lithium for 24 h and AMPK activation was determined by Western blot. Samples were probed for p-AMPK, stripped until no signal was detected and re-probed for total AMPK. The lower part of the membrane was probed against actin to normalize for protein loading. The graph shows relative protein expression of p-AMPK normalized against total AMPK and then actin. Statistical significance was analyzed by parametric t-test, *p<0.05 where n = 3 independent experiments. All graphs show mean ± SD. C, Cells were treated with either 1 mM VPA or 10 mM lithium for 24 h and IP6K1 localization was determined by fluorescence confocal microscopy. DAPI was used as nuclear staining. Scale bar = 20 μm. D, Mean fluorescence (arbitrary units, A.U.) of peripheral (cytosolic) and central (nuclear) sections of cells as shown in ‘C’. Statistical significance was analyzed by a Kolmogorov-Smirnov test, **p<0.01 where n = 3 independent experiments in which 20-30 cells were analyzed per condition. E, Western blot against MIPS protein in WT MEF cells treated with 1 mM VPA or 10 mM lithium for 24h. Graphs depict band quantification (right). Actin was used to normalize for protein loading. n = 4 independent experiments for each condition. Statistical significance was analyzed by one-way ANOVA with a Tukey *post hoc* test, *p<0.05.

Taken together, these findings support a model in which PA regulates inositol synthesis by controlling localization of the negative regulator IP6K1 (Figure 7). Increasing PA levels by activation of PLD (as a result of treatment with PMA or AMPK stimulation) results in translocation of IP6K1 to the nucleus and a decrease in MIPS protein levels.

**FIGURE 7.**
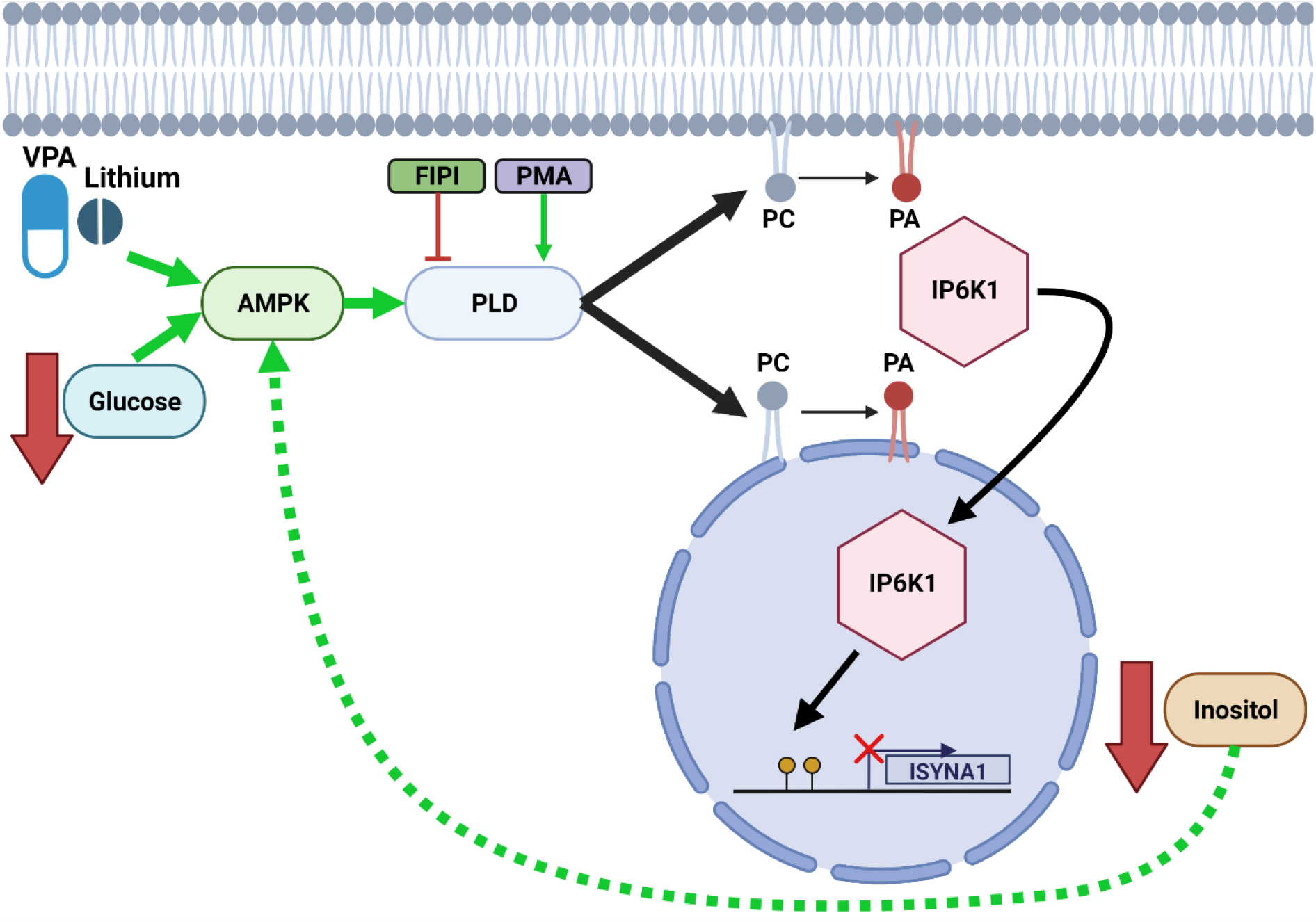
Model of PA-mediated MIPS regulation and feedback. PA can be synthesized through PLD, which converts PC into PA. PA synthesis leads to increased nuclear translocation of IP6K1 as a result of PA-IP6K1 binding. Once in the nucleus, IP6K1 induces methylation of the ISYNA1 gene promoter and inhibits MIPS expression (Yu et al., 2016). As a result, inositol levels are decreased. Inositol has been shown to negatively regulate AMPK activation (Hsu et al., 2021); therefore a decrease in inositol is predicted to lead to an increase in AMPK activity. AMPK can also be activated by low glucose, or by treatment with the mood stabilizers VPA or lithium. Activated AMPK can in turn directly activate PLD (Kim et al., 2010). Figure created with BioRender.com

## DISCUSSION

Inositol is an essential nutrient for all eukaryotic life. It serves as a precursor for phosphoinositides, inositol phosphates, and inositol pyrophosphates, which mediate a myriad of vital cellular functions related to membrane integrity, metabolic regulation, and intracellular signaling and trafficking (Case et al., 2020; Di Paolo and De Camilli, 2006; Shears, 2015; Wilson et al., 2013). Considering its importance, it is surprising that very little is known about how inositol levels are regulated in higher eukaryotes. The current study provides the first in-depth description of a mechanism for inositol regulation in mammalian cells. Accordingly, binding of plasma membrane-derived PA to the inositol pyrophosphate biosynthetic enzyme IP6K1 mediates translocation of IP6K1 to the nucleus, where it represses MIPS expression and thereby decreases inositol synthesis. IP6K1 translocation is regulated by PA. Increasing PLD-derived PA through either pharmacological treatment or physiological conditions that activate AMPK results in increased nuclear localization of IP6K and MIPS repression (Figure 7).

The exact nature of how PA influences IP6K1 localization is not clear, but the study by Yu *et al*. provided insight into this interaction (Yu et al., 2016). IP6K1 contains three functionally relevant domains that influence PA-induced nuclear translocation and MIPS repression. These include a PA-binding domain, a conserved nuclear localization signal (NLS), and an ATP-binding domain required for catalytic activity of the protein (Yu et al., 2016). Deletion of either the PA-binding domain or NLS-containing domain of IP6K1 prevents nuclear translocation and repression of MIPS. Interestingly, deletion of the ATP-binding domain has no effect on nuclear translocation but does attenuate MIPS repression. This suggests that PA-mediated MIPS repression requires both PA-IP6K1 binding and IP6K1 catalytic activity.

PA has been implicated in regulatory processes that control numerous aspects of cell growth, and specific PA species differ in their regulatory capacities (Thakur et al., 2019). For example, mono-and di-unsaturated PA species are specifically involved in recruitment and docking of secretory vesicles, whereas poly-unsaturated PA uniquely participates in fusion pore expansion during exocytosis (Tanguy et al., 2021). The site of PA synthesis dictates the specific species of PA produced (Du et al., 2004; Fazio et al., 2020; Zhukovsky et al., 2019). The finding that PA derived from the PM but not from the ER can induce nuclear translocation of IP6K1 (Figure 3) suggests that binding is dependent on the specific PA species synthesized in the PM. Interestingly, the two major isoforms of PLD differ in their subcellular locations. PLD2 is localized primarily in the PM, whereas PLD1 is primarily found in organelle membranes, such as the nuclear membrane (Tei and Baskin, 2020). Therefore, regulation of MIPS by PA may depend more on the activity of PLD2 than PLD1.

The role of IP6K1 catalytic activity in the repression of MIPS is not understood. IP6K1 catalyzes the conversion of IP_6_ to IP_7_, and IP_7_ has been shown to promote dissociation of the histone lysine demethylase JMJD2C from chromatin, resulting in increased trimethylation of histone H3 lysine 9 (H3K9me3) and transcriptional silencing of associated genes (Burton et al., 2013). Thus, IP6K1 activity may serve to silence MIPS by inducing methylation of the ISYNA1 promoter and/or increasing H3K9me3 (Burton et al., 2013).

It has been previously shown that AMPK is activated by IP_6_ and inhibited by IP_7_ (Zhu et al., 2016). Thus, the ratio of nuclear to cytosolic IP6K1 may modulate levels of IP_6_ and IP_7_ and thereby regulate AMPK activation. Interestingly, treatment of cells with the inositol-depleting drugs lithium or VPA results in AMPK activation and IP6K1 nuclear translocation (Figure 6). Lithium and VPA are known inhibitors of the inositol biosynthetic pathway; lithium uncompetitively inhibits inositol monophosphatase and VPA indirectly inhibits MIPS by an unknown mechanism (Berridge et al., 1989; Deranieh et al., 2013; Vaden et al., 2001). The current findings suggest that, in addition to these previously described mechanisms, lithium and VPA may induce inositol depletion by activating AMPK and increasing PA production. Intriguingly, a recent study demonstrated that reduced inositol levels can directly activate AMPK in MEF cells (Hsu et al., 2021). This suggests that there may be positive feedback between AMPK activation and inositol depletion wherein activated AMPK further reduces inositol levels by stimulating PLD-mediated synthesis of PA and in turn repressing MIPS expression via IP6K1 (Figure 7). Furthermore, PA can also activate AMPK through the upstream kinase LKB1, suggesting that there may be additional feedback pathways between PA levels and AMPK activation (Dogliotti et al., 2017). These scenarios are not mutually exclusive and may act in tandem to promote sustained activation of AMPK and inositol depletion.

Taken together, the current study delineates the first described mechanism for regulation of inositol synthesis in mammalian cells. This study demonstrates that transcription of the rate-limiting enzyme in the inositol biosynthetic pathway, MIPS, is negatively regulated by the abundance of a specific pool of PM-derived PA through the nuclear translocation of IP6K1 (Figure 7). This study is, to our knowledge, the first to show regulation of MIPS protein levels in mammalian cells by treatment with the mood stabilizers VPA and lithium. This novel regulatory model has important implications for the treatment of bipolar disorder and other pathological conditions using inositol-depleting mood stabilizers, as the mechanism of action of these drugs may rely on feedback between inositol depletion and AMPK activation.

## ACKNOWLEDGEMENTS

We thank the laboratory of Dr. Guangwei Du for kindly providing the GFP-PASS and RFP-PASS constructs.

## AUTHOR CONTRIBUTIONS

PL, MS, and MG conceived and designed the experiments. PL and CO performed the experiments. PL analyzed the data. PL, MS, CO and MG wrote the paper. MS highly contributed to editing and revision of the manuscript.

## ABBREVIATIONS

MEFs: mouse embryonic fibroblasts
PA: phosphatidic acid
MIPS: myo-inositol-3-phosphate synthase
IP6K1: inositol hexakisphosphate kinase
PLD: Phospholipase D
VPA: valproate
G6P: glucose-6-phosphate
AMPK: 5’ AMP-activated protein kinase
BD: bipolar disorder
LPAAT: lysophosphatidic acid acyltransferase
DAG: diacylglycerol
DGK: diacylglycerol kinase
PC: phosphatidylcholine
PMA: Phorbol 12-myristate 13-acetate
RIPA: radioimmunoprecipitation assay
PM: optoPLD
ER-optoPLD: endoplasmic reticulum-derived PA
IP7: inositol pyrophosphate 7

## DATA AVAILABILITY

All data is contained within the article.

